# Impaired cholesterol homeostasis in SNr parvalbumin neurons disrupts basal ganglia circuitry for action selection

**DOI:** 10.1101/2025.10.21.683627

**Authors:** Tomoya Ozaki, Yumi Morisugi, Mei Kamino, Hizuki Takahashi, Miki Iwata, Kazuhiro Nakamura, Honoka Maki, Michiko Shirane

**Author notes:** **Correspondence:** Michiko Shirane. These authors have contributed equally to this work and share first authorship (Tomoya Ozaki, Yumi Morisugi, Mei Kamino, Hizuki Takahashi, Miki Iwata).

## Abstract

The basal ganglia–thalamo–cortical loop regulates motor, cognitive, and emotional behaviors and plays central roles in action selection, decision-making, habit formation, and reward learning. The substantia nigra pars reticulata (SNr), a major output nucleus of the basal ganglia, exerts temporally precise control over behavior through tonic inhibitory output mediated by parvalbumin (PV)-positive GABAergic neurons.

Accumulating evidence implicates abnormalities in cholesterol metabolism in neurological disorders; however, how cholesterol metabolic dysfunction affects basal ganglia circuit regulation remains largely unknown. In our previous studies, we demonstrated brain-specific cholesteryl ester accumulation and attention-deficit/hyperactivity disorder (ADHD)-like behavioral abnormalities in mice lacking PDZD8.

Here, we show that PDZD8 deficiency is associated with selective lipofuscin-like lipid accumulation in SNr-PV neurons, accompanied by reduced SNr projections to thalamic and midbrain dopaminergic targets and increased striatonigral connectivity from caudate-putamen neurons. We propose that lipid accumulation may impair SNr-PV neuron function, potentially leading to relative disinhibition of SNr target regions and disruption of the basal ganglia–thalamo–cortical loop. These circuit abnormalities are likely to disrupt action selection and contribute to ADHD-like behavioral phenotypes.

Together, our findings identify SNr-PV neurons as a previously unrecognized site of vulnerability to cholesterol metabolic dysfunction and provide a potential mechanistic link between impaired cholesterol homeostasis, basal ganglia circuit disruption, and ADHD-like behaviors. More broadly, our findings highlight an important role for cholesterol metabolism in maintaining basal ganglia circuitry involved in action selection.

## INTRODUCTION

### SNr is a major output nucleus of the basal ganglia

The basal ganglia are a key component of the voluntary motor system and play essential roles in cognition, sensorimotor integration, emotion, and learning. Dysfunction of basal ganglia circuits has been implicated in a wide range of neurological and neuropsychiatric disorders^1–3^. The basal ganglia–thalamo–cortical loop represents a fundamental neural network motif that regulates action selection through two complementary pathways: the direct pathway (striatonigral pathway), which facilitates selected actions (“Go” pathway), and the indirect pathway (striatopallidal pathway), which suppresses competing or inappropriate actions (“No-Go” pathway). Both pathways ultimately converge on the substantia nigra pars reticulata (SNr), one of the principal output nuclei of the basal ganglia^2–4^. The SNr consists predominantly of inhibitory GABAergic neurons that project to specific thalamic nuclei and brainstem regions. SNr neurons exhibit high-frequency tonic firing, thereby maintaining downstream targets under continuous inhibitory control. Transient reductions in firing produce disinhibition of downstream circuits, enabling movement initiation and behavioral selection. Through this mechanism, the SNr exerts temporally precise control over motor and behavioral outputs during ongoing actions. Recent studies have further revealed substantial anatomical and functional heterogeneity within the SNr. In particular, its caudal and lateral subregions receive inputs from distinct striatal territories and exhibit highly organized projection patterns, suggesting specialized roles in movement control and action selection^3,5^.

### SNr dysfunction is associated with ADHD-related phenotypes

Attention-deficit/hyperactivity disorder (ADHD) is characterized by inattention, hyperactivity, and impulsivity and has been linked to dysregulated dopaminergic and noradrenergic neurotransmission^6,7^. Reduced γ-aminobutyric acid (GABA) levels have also been reported in cortical and subcortical regions of individuals with ADHD. Among inhibitory neuronal populations, parvalbumin (PV)-positive neurons play critical roles in attentional processes and motor control, and their dysfunction has been implicated in several neuropsychiatric disorders, including schizophrenia, autism spectrum disorder (ASD), and ADHD^8,9^. Within the basal ganglia, PV-positive neurons in the SNr provide powerful inhibitory output that shapes circuit activity through tonic inhibition and disinhibitory control of downstream targets. Notably, recent studies demonstrated that selective loss of histamine H₂ receptor (H₂R) signaling in SNr PV neurons reduces their activity and induces ADHD-like behavioral phenotypes, including hyperactivity, impulsivity, and inattention^10–12^. Importantly, dysfunction of SNr PV neurons, rather than medial prefrontal cortical PV neurons, was associated with elevated striatal dopamine levels, suggesting that the SNr serves as a critical node linking basal ganglia circuit dysfunction to dopaminergic dysregulation in ADHD^12^. Furthermore, ADHD frequently co-occurs with ASD and other neurodevelopmental disorders, suggesting the existence of shared pathogenic mechanisms affecting brain development and circuit function.

### Cholesterol metabolism is essential for neuronal function

Abnormal lipid accumulation has long been associated with neurodegenerative disorders, including Alzheimer’s disease (AD)^13,14^. The identification of apolipoprotein E4 (APOE4) as the strongest genetic risk factor for AD highlighted the importance of cholesterol metabolism in brain health^15^. Cholesteryl esters (CEs), neutral lipids involved in cholesterol transport and storage, accumulate abnormally in the brains of AD patients and multiple AD animal models ^16–19^. In neurons and glial cells carrying APOE4, oxidized lipids and damaged proteins accumulate within lysosomes together with CEs, leading to the formation of autofluorescent lipofuscin granules^20^. Lipofuscin accumulation has been shown to promote lysosomal stress and neuroinflammation and is thought to contribute to neuronal dysfunction and degeneration^21–23^. Although lipid dysregulation has been extensively studied in neurodegenerative diseases, its contribution to neurodevelopmental and neuropsychiatric disorders remains incompletely understood.

### PDZD8 regulates cholesterol homeostasis in the brain

We previously identified the lipid transport protein PDZD8 as a tethering factor at endoplasmic reticulum (ER)-endosome membrane contact sites (MCSs), where it contributes to endosome–lysosome maturation through cholesterol transport^24,25^. PDZD8 belongs to the TULIP (tubular lipid-binding) superfamily and contains a synaptotagmin-like mitochondrial lipid-binding protein (SMP) domain that mediates lipid transfer between membranes^25–28^. PDZD8-deficient (PDZD8^⁻/⁻^) mice exhibit brain-specific accumulation of CEs and lipofuscin-like deposits associated with lysosomal dysfunction^25,29^. Remarkably, these mice display ADHD-like behavioral abnormalities, including hyperactivity, reduced anxiety and fear responses, impairments in fear-conditioning memory and working memory, and enhanced sensorimotor gating^30^. Moreover, pathogenic PDZD8 variants in humans cause syndromic intellectual disability accompanied by regional brain overgrowth and atrophy and have been linked to intellectual developmental disorder with autism and dysmorphic features (IDDADF)^31,32^. Despite these observations, the specific brain regions and neural circuits affected by PDZD8 deficiency remain unknown.

In the present study, we identified prominent lipid accumulation in the SNr of PDZD8^⁻/⁻^ mice and found that this pathology was associated with alterations in basal ganglia–thalamo–cortical loop, including increased striatonigral connectivity. These findings suggest that lipid metabolic abnormalities in the SNr are associated with remodeling of basal ganglia circuitry and provide a potential mechanistic framework linking disrupted lipid homeostasis, circuit dysfunction, and ADHD-like behavioral phenotypes.

## RESULTS

### Marked lipid accumulation in the SNr in PDZD8^−/−^ mice

Given that PDZD8^−/−^ mice exhibit abnormal CE accumulation in the brain, we investigated the affected regions. Comprehensive staining with the neutral lipid marker LipiDye II was performed in the brains of PDZD8^+/+^ and PDZD8^−/−^ mice. As a result, marked lipid accumulation was detected in several discrete regions of PDZD8^−/−^ mice compared with PDZD8^+/+^ mice, including SNr, dorsal raphe nucleus (DR), basolateral amygdala (BLA), and central amygdala (CeA), with the most prominent abnormality observed in the SNr (Fig. 1A). Similar results were consistently confirmed in at least three independent pairs of PDZD8^+/+^ and PDZD8^−/−^ mice and were statistically ranked. Furthermore, these findings were validated not only by LipiDye II staining but also by LipidTOX staining. Both probes specifically label neutral lipids within the hydrophobic core of lipid droplets (LDs) with a high signal-to-noise ratio (Fig. 1B, C). In addition, high-resolution Z-stack imaging revealed numerous ring-shaped dense intracellular LDs within SNr cells of PDZD8^−/−^ mice, suggestive of lipofuscin-like deposits (Fig. 1D). Collectively, these results indicate that the SNr is among the brain regions showing the most prominent lipid accumulation in PDZD^−/−^ mice.

**Figure 1.**
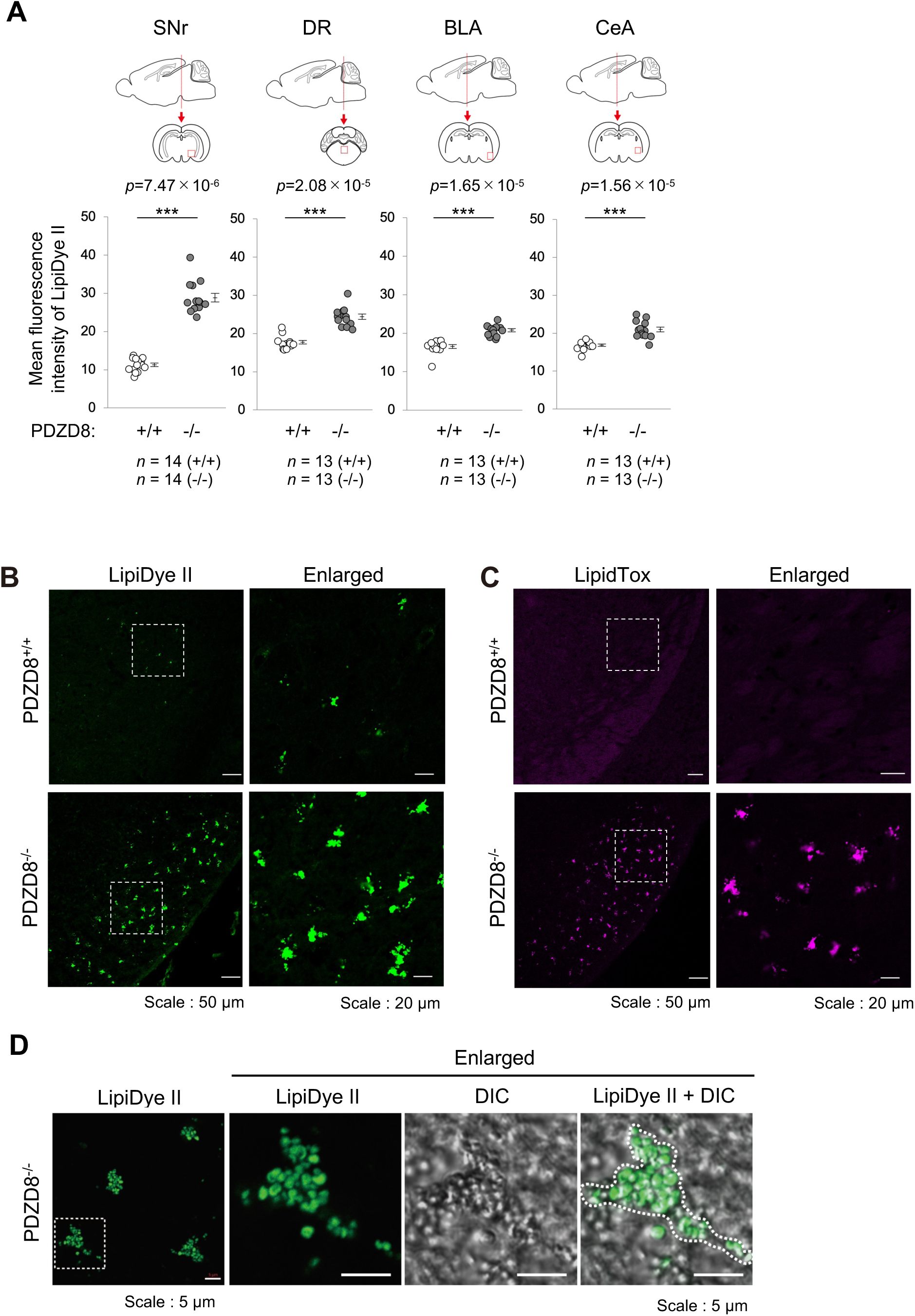
Lipofuscin-like deposits are markedly increased in the SNr in PDZD8^−/−^mice. Neutral lipid staining in PDZD8^+/+^ and PDZD8^−/−^ mouse brains. (A) Dot plots showing mean fluorescence intensity of LipiDye II in SNr, dorsal raphe (DR), basolateral amygdala (BLA), and central amygdala (CeA). Schematic brain sections indicate the regions analyzed. Red lines in sagittal sections (upper) indicate the positions of coronal sections (lower), and red boxes mark the analyzed regions. Sample size (*n*) denotes the number of images from three mice per each genotype. (B, C) Confocal images of LipiDye II (B) and LipidTox (C) in SNr. Boxed regions (left) are shown at higher magnification (right). Scale bars, 50 µm (left) and 20 µm (right). (D) High-resolution images of LipiDye II in SNr of PDZD8^−/−^ mice. Boxed regions (left) are shown enlarged (right). A differential interference contrast (DIC) image is also shown. The dotted line in merged image outlines a cell periphery densely packed with lipofuscin (right). Scale bars, 5 µm.

### PDZD8 expression in PV neurons in the SNr

To identify PDZD8-expressing cells in the SNr, immunofluorostaining using an anti-PDZD8 antibody was performed in PDZD8^+/+^ mice. PDZD8 was strongly expressed in the SNr, as well as in the ventral tegmental area (VTA) and substantia nigra pars compacta (SNc) in the midbrain (Fig. 2A). Dopamine transporter (DAT) staining was used as a marker to distinguish these regions. Within the SNr, PDZD8 was highly expressed in most parvalbumin (PV)-positive neurons (Fig. 2B). Notably, both PDZD8 and PV expression were enriched in the central-to-lateral regions of the SNr rather than being uniformly distributed throughout the entire nucleus (Supplementary Fig. 1A, B). In addition to PV neurons, PDZD8 expression was also detected in a subset of cells morphologically resembling astrocytes (Fig. 2C, arrowheads). Consistent with this observation, a portion of PDZD8 immunoreactivity colocalized with the astrocytic marker GFAP (Fig. 2D). These results indicate that the majority of PDZD8-positive cells in the SNr are PV neurons, whereas a smaller population consists of astrocytes. In contrast, PDZD8 expression was not detected in microglia or oligodendrocytes (data not shown). No obvious abnormalities were observed in the distribution or morphology of PV neurons in the midbrain regions surrounding the SNr or within the SNr itself in PDZD8^−/−^ mice in comparison to PDZD8^+/+^ (Supplementary Fig. 1C). In contrast, in the SNc and VTA, PDZD8 was expressed in DAT- and tyrosine hydroxylase (TH)-positive dopaminergic neurons (Supplementary Fig. 2A, B). These findings indicate that the identity of PDZD8-expressing cells varies among brain regions.

**Figure 2.**
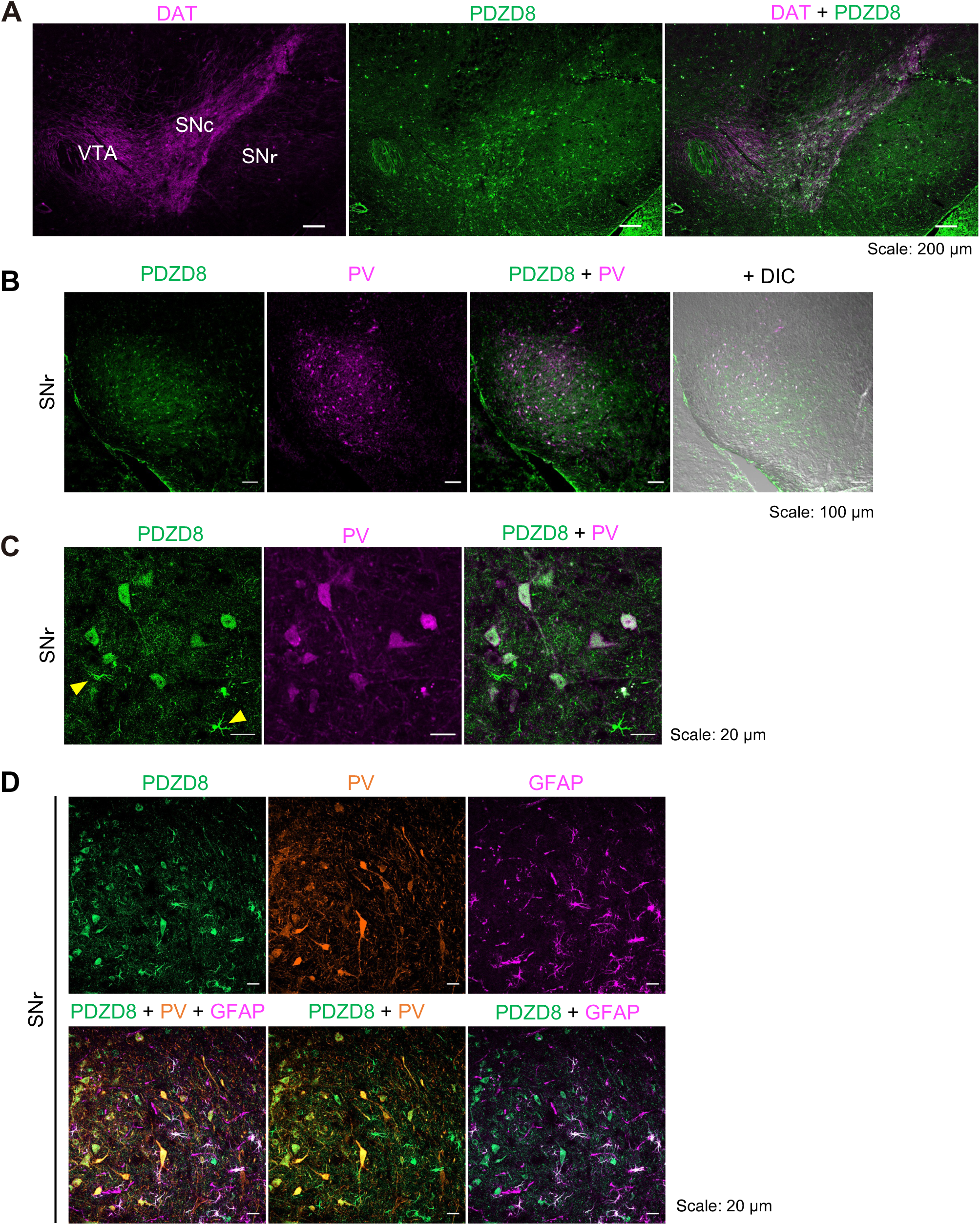
PDZD8 is highly expressed in PV neurons in the SNr. Distribution of PDZD8 in the SNr in wild-type mouse brains. (A) Confocal images of region marker dopamine transporter (DAT, magenta) and PDZD8 (green) in the midbrain. Regions corresponding to substantia nigra pars compacta (SNc), ventral tegmental area (VTA), and substantia nigra pars reticulata (SNr) are indicated. The merged image is also shown. Scale bars, 200 µm. (B) Confocal images of PDZD8 (green) and parvalbumin (PV, magenta) in SNr. Merged images witout and with DIC are also shown. Scale bars, 100 µm. (C) Enlarged images of PDZD8 (green) and PV (magenta) in (B). Astrocyte-like cells are indicated (arrowheads). Scale bars, 20 µm. (D) Confocal images of PDZD8 (green) , PV (orange), and GFAP (magenta) in SNr. The merged images are also shown. Scale bars, 20 µm.

### Lipofuscin deposits are enriched in the SNr in PDZD8^−/−^ mice

Because the SNr is a broad structure with region-specific cellular distributions, we further analyzed the spatial pattern of lipid accumulation within the SNr of PDZD8^−/−^ mice in more detail. Serial sections encompassing the SNr region from PDZD8^+/+^ and PDZD8^−/−^mice were stained with LipiDye II together with an anti-DAT antibody as a regional marker. Small punctate fluorescent signals of LipiDye II were broadly observed throughout the SNr in both PDZD8^+/+^ and PDZD8^−/−^ mice, although they tended to be more abundant in PDZD8^−/−^ mice. In contrast, intense large aggregated fluorescent signals of LipiDye II were prominently detected in the caudal SNr of PDZD8^−/−^ mice (Fig. 3A). Furthermore, in PDZD8^−/−^ mice, marked DAT fluorescent signals showing a pattern similar to that of LipiDye II were detected in the caudal slices of atlas #87–91 (Fig. 3A). These intense LipiDye II and DAT fluorescent signals were specifically localized to the lateral portion of the caudal SNr in PDZD8^−/−^ mice (Fig. 3B–D). However, DAB immunostaining using the anti-DAT antibody revealed no differences in DAT expression patterns between PDZD8^+/+^ and PDZD8^−/−^ mice (Fig. 3E), and no large aggregated DAT-positive signals were observed in the SNr region of PDZD8^−/−^ mice (Fig. 3F). In addition, no significant abnormalities in the morphology of DAT-positive neurons were detected in either the SNr or SNc regions of PDZD8^−/−^ mice by DAB staining (Supplementary Fig. 3A, B). These findings suggested that the large aggregated fluorescent signals observed in the caudal SNr of PDZD8^−/−^ mice were nonspecific. To further examine this possibility, the same region of SNr in PDZD8^−/−^ mice was subjected to triple immunofluorescence staining for LipiDye II, DAT, and GFAP. Remarkably, all three fluorescent signals, which should not biologically overlap, completely colocalized (Supplementary Fig. 3C). Therefore, the large aggregated fluorescent signals in the caudal SNr of PDZD8^−/−^ mice were consistent with autofluorescent lipofuscin-like deposits rather than specific immunoreactivity. Collectively, these results indicate that lipofuscin-like deposits preferentially accumulate in the lateral portion of the caudal SNr in PDZD8^−/−^ mice.

**Figure 3.**
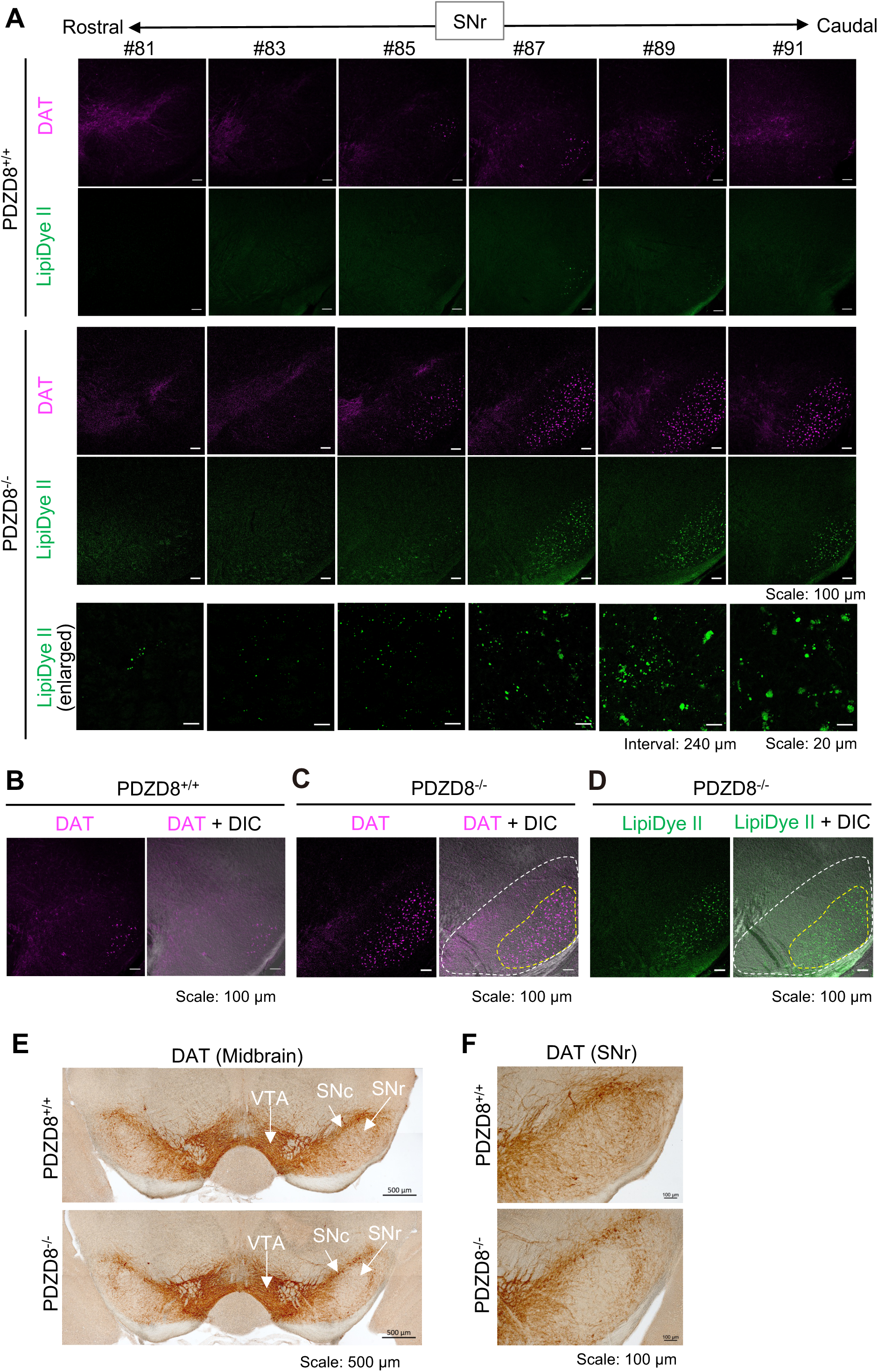
Lipofuscin-like deposits are selectively enriched in the caudal SNr of PDZD8^−/−^ mice. Images of LipiDye II and DAT in SNr in PDZD8^+/+^ and PDZD8^−/−^ mouse brains. (A) Confocal images of DAT (magenta) and LipiDye II (green) in serial sections of SNr from rostral (left) to caudal (right) regions at 240 µm intervals. Scale bars, 100 µm. For PDZD8-/-, enlarged images of LipiDye II are also shown (lower). Scale bars, 20 µm. Section numbers corresponding to Allen mouse brain atlas image are indicated (top). (B, C) Confocal images of DAT (magenta) in the SNr of the section #87 in (A) of PDZD8^+/+^ (B) and PDZD8^−/−^ (C) mouse brains. The merged images with DIC are also shown. Scale bars, 100 µm. (D) Confocal images of LipiDye II (green) in the adjacent section to (C) in PDZD8^−/−^mouse brain. The merged images with DIC are also shown. The regions corresponding to the SNr or enriched in lipofuscin-like deposits are outlined with white or yellow dotted lines, respectively (C, D). Scale bars, 100 µm. (E) DAB images of DAT in midbrain in PDZD8^+/+^ and PDZD8^−/−^. The region of VTA, SNc, and SNr are indicated with arrows. Scale bars, 500 µm. (F) High-resolution images of DAT in SNr in (E). Scale bars, 100 µm.

### Lipofuscin deposits in PV neurons of the caudal SNr in PDZD8^−/−^ mice

In the caudal SNr, where lipofuscin accumulation was observed, PDZD8 was predominantly expressed in PV neurons and, to a lesser extent, in astrocytes (Fig. 2). However, these two cell populations exhibited distinct spatial distributions: PV neurons were concentrated in the lateral region, whereas astrocytes were distributed throughout the SNr (Fig. 4A–C). To determine which cell type contributed to lipid accumulation in the caudal SNr of PDZD8^−/−^ mice, we performed Z-stack three-dimensional imaging of LipiDye II and GFAP immunofluorostaining. The pattern of lipid deposition differed markedly between the medial and lateral regions of the caudal SNr. In the medial region, lipid accumulation was primarily observed as small extracellular puncta, potentially representing extracellular lipoprotein deposits. In contrast, the lateral region contained not only these extracellular puncta but also large intracellular deposits resembling lipofuscin (Fig. 4D, E). At higher magnification, these lipofuscin-like intracellular deposits consisted of clusters of small ring-shaped structures similar to those observed in Fig. 1D. Three-dimensional reconstruction enabled clear discrimination between intracellular and extracellular LipiDye II signals (Supplementary movies A, B). Notably, two types of intracellular lipid deposits were identified in the lateral SNr: deposits localized within astrocytic processes (white arrowheads) and deposits that densely filled the cytoplasm of neuronal cell bodies (yellow arrows). Furthermore, co-fluorostaining with LipiDye II and anti-PV antibodies revealed lipofuscin-like structures within a subset of PV neurons (Fig. 4F). Notably, many of the cells containing deposits appeared morphologically atrophied and weakened compared to cells without deposites (Fig. 4F, Supplementary Fig. 4A–D). Together, these findings suggest that lipid accumulation in the SNr includes at least two components: extracellular lipoprotein-like deposits distributed throughout the caudal SNr and intracellular lipofuscin-like deposits observed in PV neurons and astrocytes in the lateral caudal SNr.

**Figure 4.**
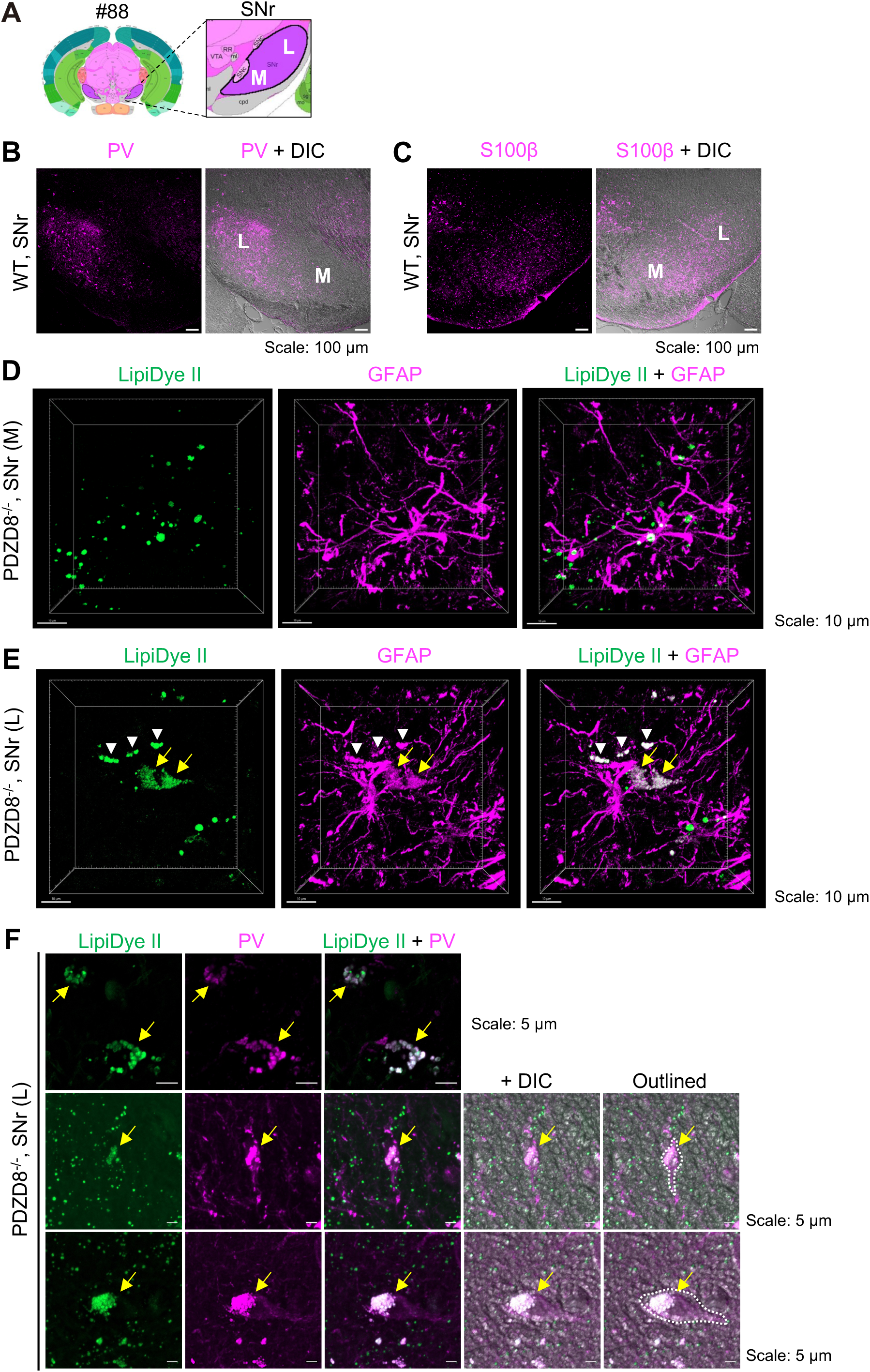
Lipofuscin deposits in PV neurons in the lateral SNr in PDZD8−/− mice. (A) Image of coronal section from Allen mouse brain atlas #88 (left) and enlarged SNr (right) with the medial (M) and lateral (L) regions indicated. (B, C) Confocal images of PV (B) or S100β in SNr of section #88 in wild-type mouse brain. The merged images with DIC are also shown. Scale bars, 100 µm. (D, E) 3D images of LipiDye II (green) and GFAP (magenta) in medial (D) and lateral (E) regions of caudal SNr of section #88 in PDZD8^−/−^ mice, which are reconstructed from z-stack confocal images using Imaris software. LipiDye II-positive structures in astrocytic processes (white arrowheads) and neural soma (yellow arrows) are indicated. Scale bars, 10 µm. (F) Confocal images of LipiDye II (green) and PV (magenta) in PV neurons containing lipofuscin deposits in lateral regions of caudal SNr in PDZD8^−/−^ mice. LipiDye II-positive structures in PV neurons (yellow arrows) are indicated. Scale bars, 5 µm.

### Decreased projection from the SNr in PDZD8^−/−^ mice

To determine whether lipofuscin accumulation in SNr PV neurons alters SNr-associated neural circuits, we compared the projection patterns of the SNr between PDZD8^+/+^ and PDZD8^−/−^ mice. Because the SNr serves as a major output nucleus of the basal ganglia, the anterograde tracer palmitoylated GFP (palGFP) was injected into the SNr to visualize its efferent projections. The tracer injection sites were comparable between the two genotypes (Fig. 5A, B). Compared with PDZD8^+/+^ mice, PDZD8^−/−^ mice exhibited a marked reduction in SNr projections to the ventral anterior lateral/medial thalamic nuclei (VAL/VM) and the nigrostriatal tract (nst) (Fig. 5C), as well as to the SNc and VTA (Fig. 5D). Quantitative analysis confirmed significant decreases in projection density in these target regions (Fig. 5E), suggesting reduced SNr connectivity to these target regions in PDZD8^−/−^ mice (Fig. 5F). Similar results were reproduced in at least two independent cohorts of PDZD8^+/+^ and PDZD8^−/−^ mice, and the differences were statistically significant (Supplementary Fig. 5A–D). These findings indicate altered SNr projection patterns in PDZD8^−/−^ mice. Given that SNr neurons normally exert tonic inhibitory control over their downstream targets, the reduced SNr projections observed in PDZD8^−/−^ mice may be consistent with reduced inhibitory influence of the SNr on downstream targets.

**Figure 5.**
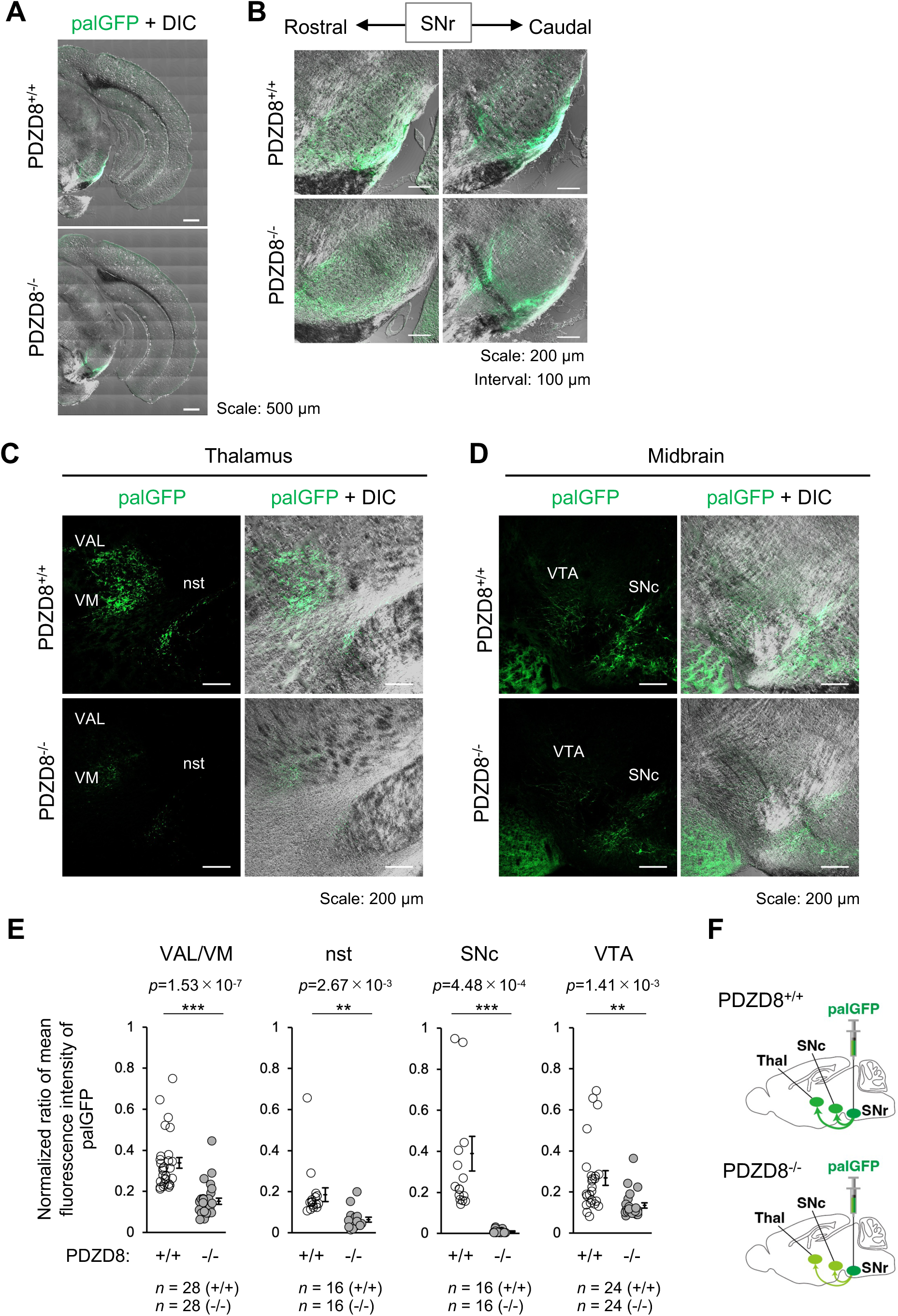
Decreased projection from the SNr in PDZD8^−/−^ mice. (A, B) Confocal images showing palmitoylated GFP (palGFP) injection sites in SNr merged with DIC images. Tiling images of cerebral hemisphere (A) and serial sections from rostral (left) to caudal (right) regions at 100 µm intervals focusing on SNr (B). Scale bars, 500 µm (A); 200 µm (B). (C) Confocal palGFP images merged with DIC of thalamic regions showing SNr projection targets. PalGFP signals indicate SNr projections to the ventral anterior-lateral complex of the thalamus (VAL), ventromedial thalamus (VM), and the nigrostriatal tract (nst). Scale bars, 200 µm. (D) Confocal palGFP images merged with DIC of midbrain regions showing SNr projections to the substantia nigra pars compacta (SNc) and ventral tegmental area (VTA). Scale bars, 200 µm. (E) Dot plots showing normalized ratio of mean fluorescence intensity of palGFP at VAL/VM, nst, SNc, and VTA. Sample size (*n*) denotes the number of images from two mice per each genotype. (F) Schematic illustration summarizing the anterograde tracing results. Projections from the SNr to VM/nst and SNc/VTA are reduced in PDZD8⁻/⁻ mice compared with PDZD8^⁺/⁺^ mice. Injection sites of palGFP in the SNr are indicated.

### Increased striatal projection to the SNr in PDZD8^−/−^ mice

We next examined neurons projecting to the SNr in PDZD8^+/+^ and PDZD8^−/−^ mice. Retrograde tracing was performed by injecting Alexa594-conjugated cholera toxin subunit B (Alexa594-CTb) into the SNr. The injection sites were comparable between the two genotypes (Fig. 6A, B). Compared with PDZD8^+/+^ mice, PDZD8^−/−^ mice exhibited a marked increase in retrogradely labeled neurons in the caudate-putamen (CP), indicating an increased number of retrogradely labeled neurons in the CP (Fig. 6C–E). This finding was independently reproduced in an additional pair of PDZD8^+/+^ and PDZD8^−/−^ mice using Alexa488-conjugated CTb (Alexa488-CTb), confirming the robustness of the result (Supplementary Fig. 6A–E; Fig. 6F). The increased CP-to-SNr projection observed in PDZD8^−/−^ mice suggests altered striatonigral connectivity. Consistent with this interpretation, increased CTb labeling was detected in neurons surrounded by dopamine D1 receptor (D1R) immunoreactivity in PDZD8^−/−^ mice, further supporting the possibility that D1R-positive striatonigral neurons contribute to the observed increase in SNr-projecting neurons in these animals (Fig. 6H).

**Figure 6.**
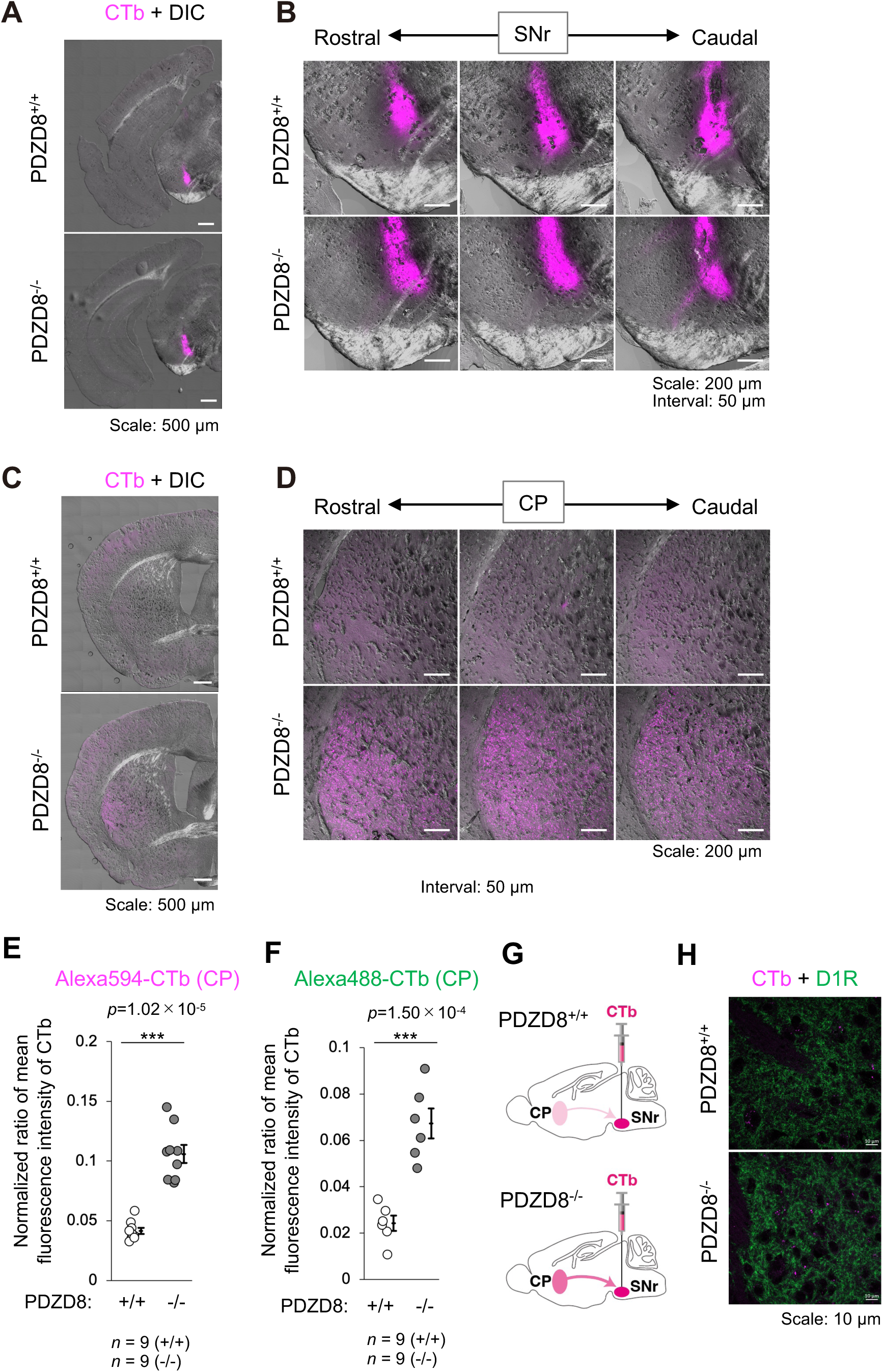
Increased striatal projection to the SNr in PDZD8^−/−^ mice. (A, B) Confocal images of Alexa594-conjugated cholera toxin subunit B (Alexa594-CTb) injected to SNr merged with DIC images. Tiling images of cerebral hemisphere (A) and serial sections from rostral (left) to caudal (right) regions at 50 µm intervals focusing on SNr (B). Scale bars, 500 µm (A) and 200 µm (B). (C, D) Confocal images of Alexa594-CTb merged with DIC images of brain region containing caudate-putamen (CP) in striatum, indicating projection sources. Tiling images of cerebral hemisphere (C) and serial sections from rostral (left) to caudal (right) regions at 50 µm intervals focusing on CP (D). Scale bars, 500 µm (C) and 200 µm (D). (E) Confocal images of Alexa594-CTb merged with D1R images of CP in (D). (F, G) Dot plots showing normalized ratio of mean fluorescence intensity of Alexa594-CTb (F) or Alexa488-CTb (G) at CP. Sample size (*n*) denotes the number of images from one mouse per each genotype. (H) Schematic representation of striatal projections to SNr, showing increased projections in PDZD8^−/−^ mice compared with PDZD8^+/+^ mice. Injection sites of CTb to SNr are also indicated.

### Enhanced striatonigral innervation in PDZD8^−/−^ mice

Because the retrograde tracing experiments suggested enhanced input from the CP to the SNr in PDZD8^−/−^ mice, we next examined medium spiny neurons (MSNs), the DARPP-32-positive principal neurons that give rise to the striatonigral direct pathway. In the CP of PDZD8^+/+^ mice, PDZD8 was strongly expressed in DARPP-32-positive neurons (Fig. 7A). We then assessed DARPP-32 immunoreactivity in the SNr, a major target of striatonigral projections. Compared with PDZD8^+/+^ mice, PDZD8^−/−^ mice exhibited a significant increase in DARPP-32 immunoreactivity within the SNr (Fig. 7B). This finding was further confirmed in sagittal brain sections, which also showed significantly elevated DARPP-32 immunoreactivity in the SNr of PDZD8^−/−^ mice relative to PDZD8^+/+^ controls (Fig. 7C). Together, these findings suggest increased striatonigral innervation of the SNr in PDZD8^−/−^ mice and are consistent with altered activity of the striatonigral direct pathway.

**Figure 7.**
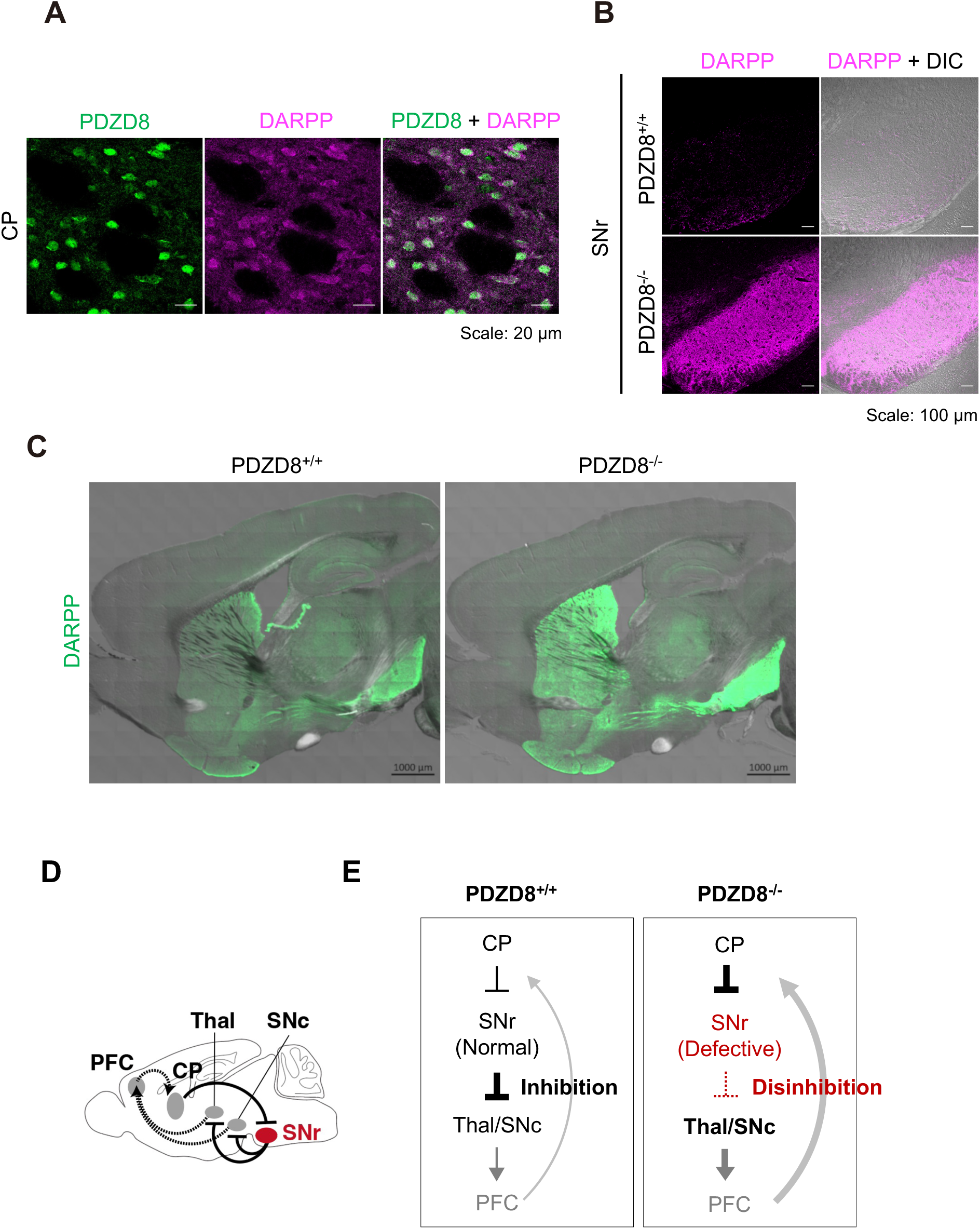
Altered basal ganglia circuitry in PDZD8^−/−^ mice. (A) Confocal images of PDZD8 (green) and DARPP-32 (magenta) in CP of wild-type mice, with merged image. Scale bars, 20 µm. (B) Confocal images of DARPP-32 and merged with DIC in SNr of PDZD8^+/+^ and PDZD8^−/−^. Scale bars, 100 µm. (C) Confocal images of DARPP-32 merged with DIC images of sagital sections of brain containing CP and SNr in PDZD8^+/+^ and PDZD8^−/−^. Scale bars, 1,000 µm. (D) Schematic representation of basal ganglia-thalamo-cortical loop. (E) Proposed model illustrating the consequences of lipid accumulation in the SNr of PDZD8^⁻/⁻^ mice. In PDZD8^⁺/⁺^ mice (left), behavior is constantly suppressed by sustained inhibitory output from PV neurons in the SNr. In contrast, in PDZD8⁻/⁻ mice, lipid accumulation in SNr-PV neurons is associated with impaired neuronal function, leading to a reduction in inhibitory output from the SNr to the VAL/VM, nst, and SNc/VTA. This leads to disinhibition of the thalamus and dopaminergic systems, and feedback via the prefrontal cortex (PFC) enhances GABAergic projections from the CP to the SNr. In PDZD8⁻/⁻ mice, disruption of this basal ganglia–thalamus–cortex circuit may contribute to dysregulation of action selection and ADHD-like behavioral phenotypes.

## DISCUSSION

### A working model linking lipid accumulation in SNr-PV neurons to basal ganglia circuit disinhibition and remodeling

Taken together, our findings support a model in which basal ganglia circuitry is disrupted in PDZD8^−/−^ mice (Fig. 7D, E). In PDZD8^+/+^ mice, PV-positive GABAergic neurons in the SNr provide tonic inhibitory output to downstream targets, thereby contributing to the suppression and appropriate selection of behavioral responses. In contrast, in PDZD8^−/−^mice, lipid accumulation within SNr-PV neurons may impair their function, potentially resulting in reduced inhibitory influence of the SNr on downstream targets. This may lead to relative disinhibition of thalamic and dopaminergic circuits, potentially altering activity within the basal ganglia–thalamocortical network. We speculate that feedback through cortical circuits, potentially involving the PFC, may contribute to the increased CP-to-SNr connectivity observed in PDZD8^−/−^ mice. Consistent with this model, PDZD8^−/−^ mice exhibited increased CP-to-SNr connectivity and elevated DARPP-32 immunoreactivity within the SNr. Collectively, these alterations are likely to disrupt the normal regulation of action selection by the basal ganglia–thalamocortical circuit. We therefore propose that dysfunction of SNr-PV neurons associated with lipid accumulation may lead to circuit-level disinhibition and remodeling, ultimately contributing to the ADHD-like behavioral abnormalities observed in PDZD8^−/−^ mice (Fig. 7D, E).

### Lipid metabolism– associated SNr disinhibition as a potential mechanism of ADHD-like behavioral abnormalities

Our findings extend previous evidence implicating SNr dysfunction in the pathophysiology of ADHD. Previous studies have demonstrated that loss of histamine H₂ receptor (H₂R) signaling in SNr PV neurons reduces inhibitory output from the SNr and induces hyperactivity and impulsive behaviors. Similarly, reduced GABAergic tone and enhanced dopaminergic transmission have been reported in ADHD patients. In the present study, PDZD8 deficiency resulted in lipid accumulation within SNr PV neurons, accompanied by reduced SNr projections to downstream target regions, enhanced striatonigral connectivity, and ADHD-like behavioral abnormalities. Accumulation of oxidized lipids within lipofuscin deposits could disrupt synaptic vesicle dynamics, membrane excitability, and ion channel stability. These findings raise the possibility that disruption of lipid homeostasis may contribute to circuit-level disinhibition within the basal ganglia network. Importantly, although the molecular causes differ, both H₂R deficiency and PDZD8 deficiency may converge on a common functional consequence: reduced inhibitory control exerted by SNr PV neurons. We therefore propose that SNr disinhibition may represent a shared circuit mechanism through which diverse cellular abnormalities impair behavioral control. More broadly, our findings raise the possibility that lipid stress in SNr neurons contributes to basal ganglia circuit dysfunction and may represent a previously underappreciated pathway involved in the development of ADHD-like phenotypes.

### Implications for neurodevelopmental and neurodegenerative disorders

Lipofuscin accumulation caused by lysosomal dysfunction is a well-established feature of neurodegenerative disorders, including Alzheimer’s disease and age-related cognitive decline. Although neurodevelopmental and neurodegenerative disorders arise at different stages of life, our findings suggest that impaired lysosomal function and disrupted cholesterol metabolism may contribute to both. In the present study, PDZD8 deficiency caused lipid accumulation, circuit dysfunction, and behavioral abnormalities, indicating that defective intracellular lipid processing can influence neural circuit function. Together with previous studies, our findings suggest that cholesterol metabolic abnormalities resulting from impaired lysosomal homeostasis may represent a shared pathogenic mechanism linking disorders of brain development and brain aging. Understanding how neuronal lipid metabolism shapes circuit function may therefore provide a unifying framework for brain disorders across the lifespan.

### Limitation of the study

The causal relationship between lipid accumulation and basal ganglia circuit dysfunction in PDZD8⁻/⁻ mice, as well as the underlying molecular mechanisms, remains to be established. Previous studies have shown that excessive lipofuscin accumulation can promote reactive oxygen species (ROS) production through lipid peroxidation, leading to lysosomal stress, chronic neuroinflammation, and progressive neuronal dysfunction^20–23^. However, whether these mechanisms contribute to the cellular and circuit abnormalities observed in PDZD8⁻/⁻ mice remains unknown. Future studies should directly quantify intracellular lipid peroxidation and ROS levels to determine the contribution of oxidative lipid stress to SNr-PV neuron dysfunction.

Although PDZD8 is broadly expressed throughout the brain, lipid accumulation in PDZD8⁻/⁻ mice was particularly prominent in the SNr. The mechanisms underlying this selective regional vulnerability remain unclear. Future studies will be required to determine whether the unique physiological properties of SNr-PV neurons, including their high-frequency firing activity and sustained inhibitory output, confer increased susceptibility to impaired cholesterol homeostasis. In addition, region-specific molecular factors, such as PDZD8-associated protein complexes or local metabolic demands, may contribute to this phenotype.

Finally, our study does not directly demonstrate that lipid accumulation is sufficient to cause the observed circuit and behavioral abnormalities. Pharmacological or genetic interventions that restore cholesterol homeostasis will therefore be important to determine whether normalization of lipid metabolism can rescue SNr function, basal ganglia circuit integrity, and action-selection behavior.

## METHODS

### Animals

Mice were housed under 12-h light/dark cycle with food and water ad libitum. All mouse experiments were approved by the animal ethics committee of Nagoya City University. All experiments were performed using at least two sets of paired mice (PDZD8^+/+^ and PDZD8^−/−^). Mouse sex and age were matched across experiments, and both mice in each pair were analyzed simultaneously. In all experiments, mice were aged 4–9 months and were either male or female. Figures show representative images from one of the multiple experiments performed. A summary table listing these mouse details is shown in Supplementary table 1.

### Mutant mice

Generation of PDZD8-KO mice was described previously^29^. In brief, Cas9 nickase mRNA, Generic tracrRNA, and target-specific crRNA were mixed and injected into C57BL/6J mouse zygotes. The resultant mutant mice were backcrossed with C57BL/6J mice before experiments. Animals were genotyped by PCR analysis.

### Antibodies and reagents

Rabbit polyclonal antibodies to PDZD8 were from Sigma-Aldrich; mouse monoclonal antibodies to TH were from Sigma-Aldrich; mouse monoclonal antibodies to GFAP were from Proteintech; guinea pig polyclonal antibodies to DAT, PV, and DARPP were from Frontier institute; Alexa Fluor 488 goat anti-rabbit IgG, Alexa Fluor 555 goat anti-mouse IgG, Alexa Fluor 555 goat anti-guinea pig IgG, Alexa Fluor 647 goat anti-guinea pig IgG, Alexa Fluor 488 cholera toxin B, Alexa Fluor 594 cholera toxin B, and LipidTox were from Thermo Fisher Scientific; LipiDye II was from Funakoshi; HRP-conjugated anti-rabbit and -mouse IgG secondary antibodies were from Promega.

### Immunostaining analysis

Mice were transcardially perfused with 2% paraformaldehyde (PFA) in phosphate-buffered saline (PBS, Wako), and the brains were dissectedfollowed by post-fixation in the same fixative. The brains were then cryoprotected by sequential immersion in 15% and 30% sucrose in PBS, embedded in OCT compound (SAKURA), and sectioned into 20-50 µm thick cryosections using a cryostat (Leica). Thin sections were prepared with a cryostat microtome (CM1850 UV, Leica). For immunofluorescence analysis, sections were incubated with primary antibodies followed by Alexa Fluor–labeled secondary antibodies in PBS containing 0.5% Triton X-100 (Tx-PBS), or stained with LipiDye II or LipidTox in PBS. The sections were observed with an LSM800 confocal microscope (Zeiss), and the images were processed for calculation of fluorescence intensity with ZEN imaging software (Zeiss). For immunohistochemistry analysis, sections were incubated with 0.3% hydrogen peroxide (H₂O₂) in PBS to block endogenous peroxidase activity, followed by primary antibodies and HRP-conjugated anti-rabbit or -mouse IgG secondary antibodies in 0.5% Tx-PBS, then visualized using a DAB staining kit (Bioenno Tech). The sections were observed with an Axio Imager A1 microscope (Zeiss). Brain regions were referenced from Allen Brain Atlas (http://atlas.brain-map.org).

### Stereotaxic injection

Mice were anesthetized intraperitoneally with a mixture of midazolam (4 mg kg^−1^, SANDOZ, Tokyo, Japan), medetomidine (0.75 mg kg^−1^, Kyoritsu Seiyaku, Tokyo, Japan) and butorphanol (5 mg kg^−1^, Meiji Seika Pharma, Tokyo, Japan) and placed in a stereotaxic frame (NARISHIGE, Tokyo, Japan). Body temperature was maintained at 37 °C. The scalp was sterilized and incised, and a small craniotomy was made above the target. Alexa Fluor 594– or 488–conjugated cholera toxin subunit B (CTb, Thermo Fisher Scientific, MA, USA) was prepared at 2 µg µL^−1^ in sterile PBS. AAV2/1-CMV-palGFP virus particles were from Dr. Nakamura^33^. 200 nl of Alexa Fluor 594-CTb was injected into the SNr (coordinates from Bregma: AP: -3.2 mm, ML: ±1.4 mm, DV: -4.4 mm). 150 nl of Alexa Fluor 488-CTb was injected into the SNr (coordinates from Bregma: AP: -3.7 mm, ML: ±1.8 mm, DV: -4.5 mm). 100 nl of AAV solution was injected into the SNr (coordinates from Bregma: AP: -3.7 mm, ML: ±1.8 mm, DV: -4.5 mm). Tracer (200 nL per site) was delivered using a Hamilton microsyringe (1701RN Neuros Syringe, 33G; Hamilton Company, NV, USA) connected to a motorized stereotaxic microinjector (NARISHIGE, Tokyo, Japan ; 100 nL min^−1^). After infusion, the needle was left in place for 10–15 min to minimize reflux before being withdrawn slowly. For each mouse, injections were made at the following stereotaxic coordinates relative to bregma: [AP -3.2], [ML ±1.4], [DV -4.4] (mm). The scalp was closed with sutures, and mice were monitored until full recovery. After surgery, mice received postoperative care including an intraperitoneal injection of antibiotic (amoxicillin, Kyoritsu Seiyaku, Tokyo, Japan; 50 mg kg^−1^, i.p.). Information of tracer injection was summarized in Supplementary table 2.

### Histological analysis of neural tracing

Mouse brain was fixed with 2% paraformaldehyde in PBS, exposed consecutively to 15% and 30% sucrose in PBS for 2 days, and embedded in OCT compound. Thin sections (50 µm) were prepared with a cryostat microtome. The sections were observed with an LSM800 confocal microscope. The distribution of the tracer at the injection site was verified, and only sections without substantial displacement of the injection site were included in the analysis. The sections were imaged using a LSM800 to acquire fluorescence images. The images were processed for calculation of fluorescence intensity with ZEN imaging software. Regions of interest (ROIs) were defined on the acquired images, and fluorescence intensity values were extracted. Background fluorescence was subtracted to obtain tracer-specific signal intensities. Tracer signals were quantified for both the injection sites and the target regions, and values from the target regions were normalized to those of the corresponding injection sites.

### Statistical analysis

For lipid imaging, statistical analysis was performed with the use of BellCurve for Excel software (Social Survey Research Information, Tokyo, Japan). Normality of data was assessed with the Shapiro-Wilk test. If the normality was not met, the Mann-Whitney U test was applied (Fig. 2A, SNr; DR; BLA). If data were normally distributed and variance was homogeneous between genotypes, comparisons were performed with Student’s *t* test. If homogeneity of variance was not assumed, Welch’s *t* test was applied instead of Student’s *t* test (Fig. 2A, CeA). For neuronal tracing, statistical analysis was performed using Microsoft Excel. Variance homogeneity was assessed using an F-test, and based on the results, appropriate t-tests, Welch’s t test (Fig. 6E, VAL/VM; SNc) or Student’s t test (Fig. 5E, Str; Fig. 6E, VTA; SNc) were applied to evaluate statistical significance. Quantitative data are presented as mean ± SE. Statistical significance of a *P* value is indicated as *p* < 0.05 (*) and *p* < 0.001 (***). Sample size (*n*) are in the figure legends.

## Acknowledgements

We thank I. Yamahata and I. Ogino for Immunofluorescent analysis, N. Takahashi, Y. Isomura, and M. Ishikawa for discussion, and the Research Equipment Sharing Center at Nagoya City University for assistance.

## Author contributions statement

M.S. conceptualized and supervised the study, acquired funding, designed experiments, and wrote the manuscript; T.O., Y.M., M.K., H.T., and M.I. performed experiments; H.M. prepared resources; K.N. provided palGFP virus particles.

## Funding information

This work was supported in part by KAKENHI grants from Japan Society for the Promotion of Science (JSPS) to M.S. (24K02122).

## Ethics

The animal study was approved by Nagoya City University, Animal Research Committee. The study was conducted in accordance with the local legislation and institutional requirements.

## Data availability and sources

The original contributions presented in the study are included in the article/ supplementary material, further inquiries can be directed to the corresponding author.

## SUPPLEMENTARY FIGURE LEGENDS

**Supplementary figure 1.**
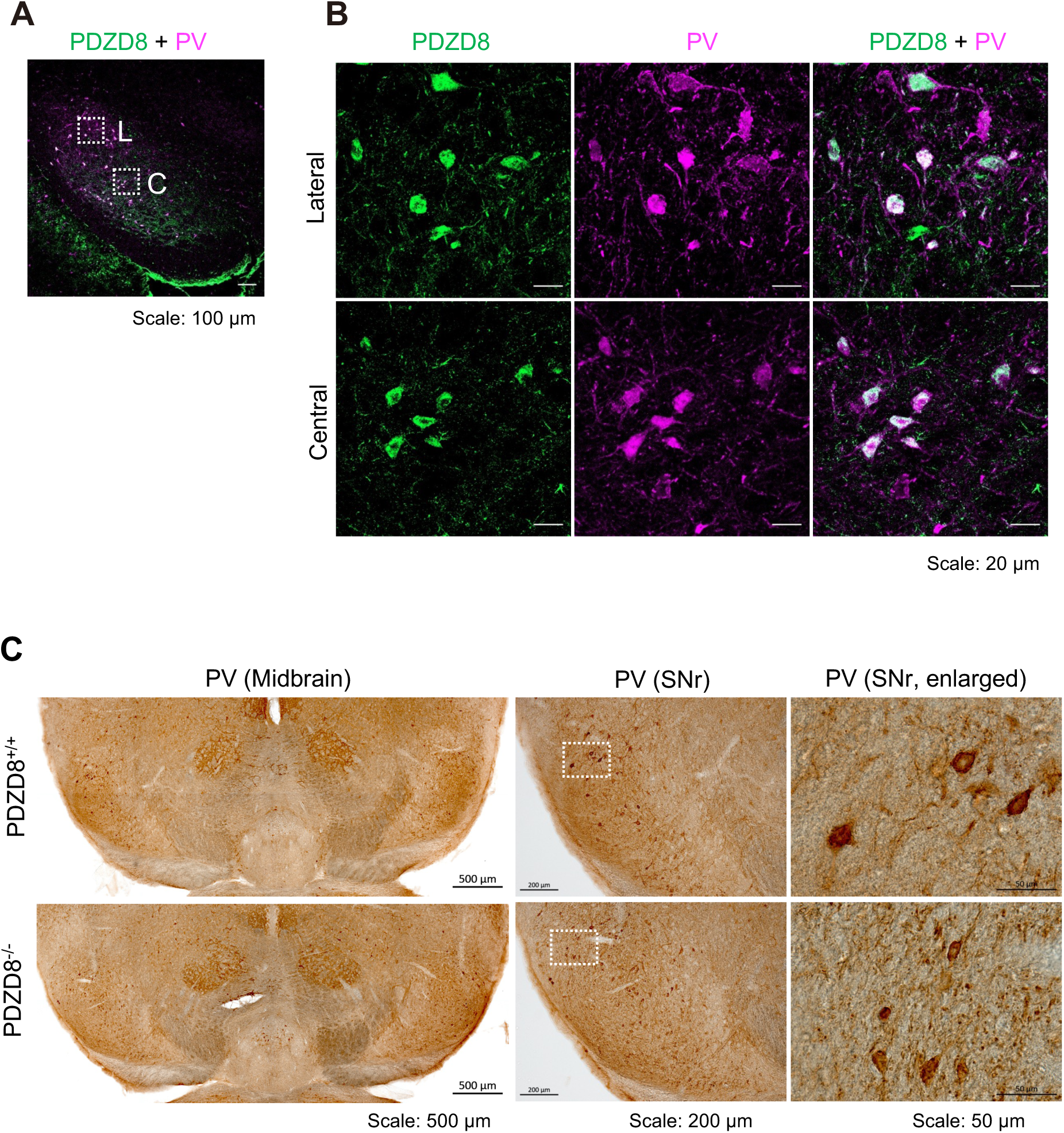
Expression of PDZD8 and PV in the SNr. (A) Confocal images of PDZD8 (green) and parvalbumin (PV, magenta) in SNr in wild-type mouse brains. The merged image is shown (A). Scale bars, 100 µm. (B) Enlarged images of lateral (B) and central (C) region in (A). Scale bars, 20 µm. (C) DAB images of PV in the midbrain (left), SNr (middle), and enlarged images of the boxed regions (right) in PDZD8^+/+^ and PDZD8^−/−^. Scale bars, 500, 200, and 50 µm, respectively.

**Supplementary figure 2.**
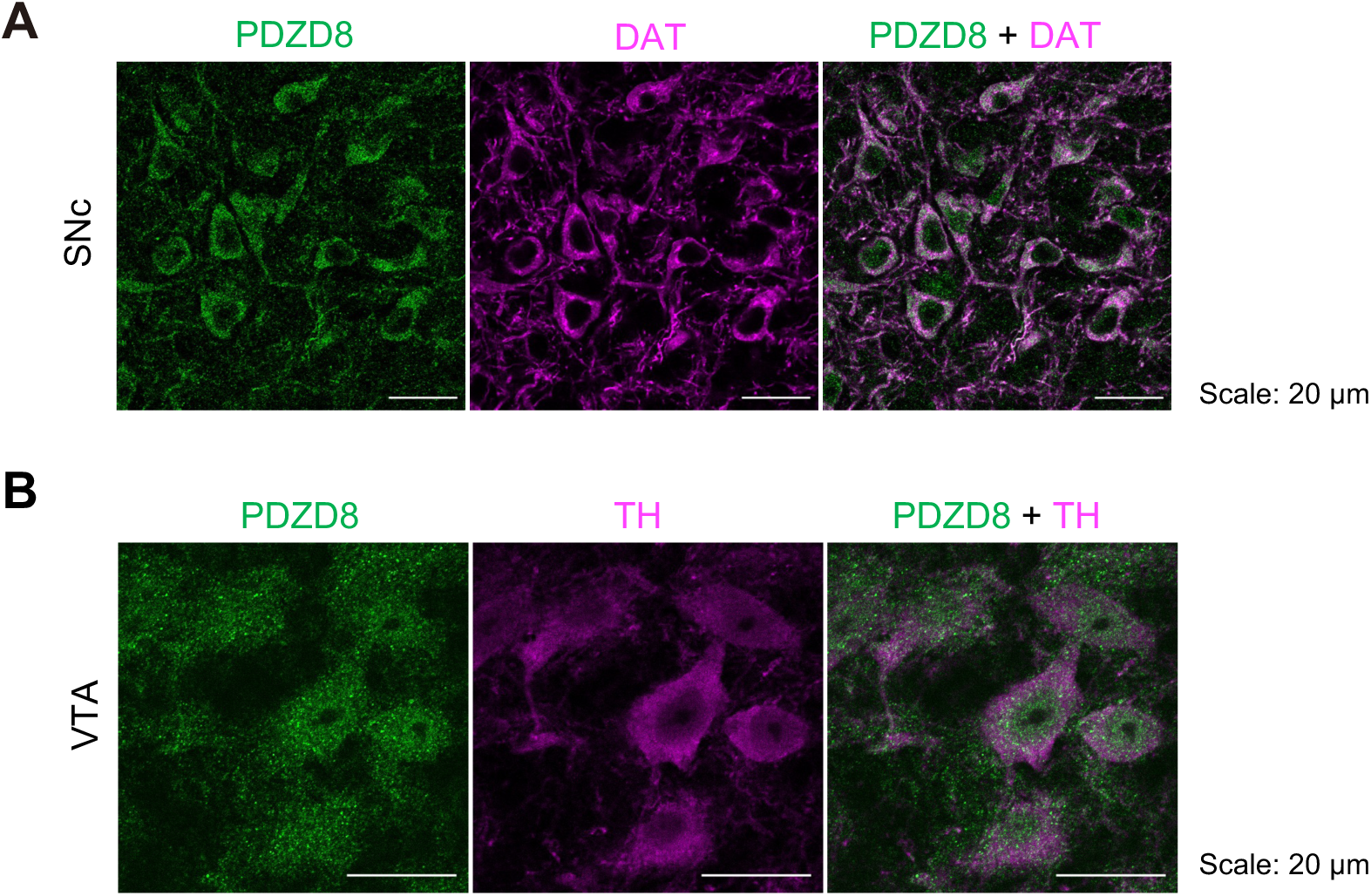
Expression of PDZD8 in PV neurons in the SNr. Immunofluorescence analysis of PDZD8 in wild-type mouse brain. (A) PDZD8 (green) and dopamine transporter (DAT, magenta) in the region corresponding to substantia nigra pars compacta (SNc). The merged image is also shown. Scale bars, 20 µm. (B) PDZD8 (green) and tyrosine hydroxylase (TH, magenta) in the region corresponding to ventral tegmental area (VTA). The merged image is also shown. Scale bars, 20 µm.

**Supplementary figure 3.**
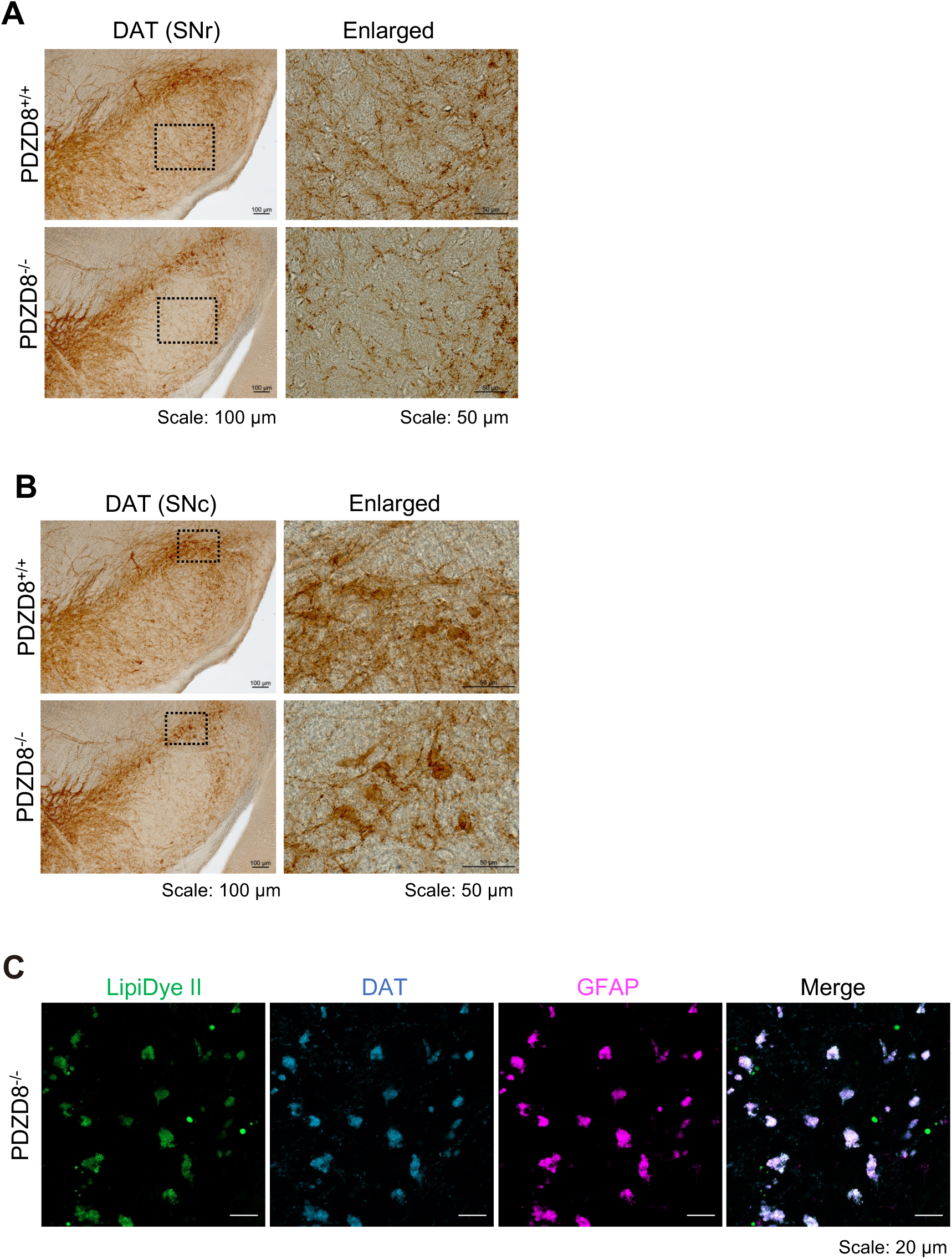
Lipofuscin in SNr does not affect DAT expression in SNr and SNc in PDZD8^−/−^ mice. (A) DAB images of DAT in SNr in PDZD8^+/+^ and PDZD8^−/−^ with enlarged images. Scale bars, 100 and 50 µm, respectively. (B) DAB images of DAT in SNc in PDZD8^+/+^ and PDZD8^−/−^ with enlarged images. Scale bars, 100 and 50 µm, respectively. (C) Lipofuscin-like deposits exhibit autofluorescence in SNr in PDZD8^−/−^ mouse brains. Confocal images of LipiDye II (green), DAT (blue), and GFAP (magenta) in the SNr in PDZD8^−/−^ mouse brain corresponding to the region with a lot of lipofuscin deposits. The merged image is also shown. Scale bars, 20 µm.

**Supplementary figure 4.**
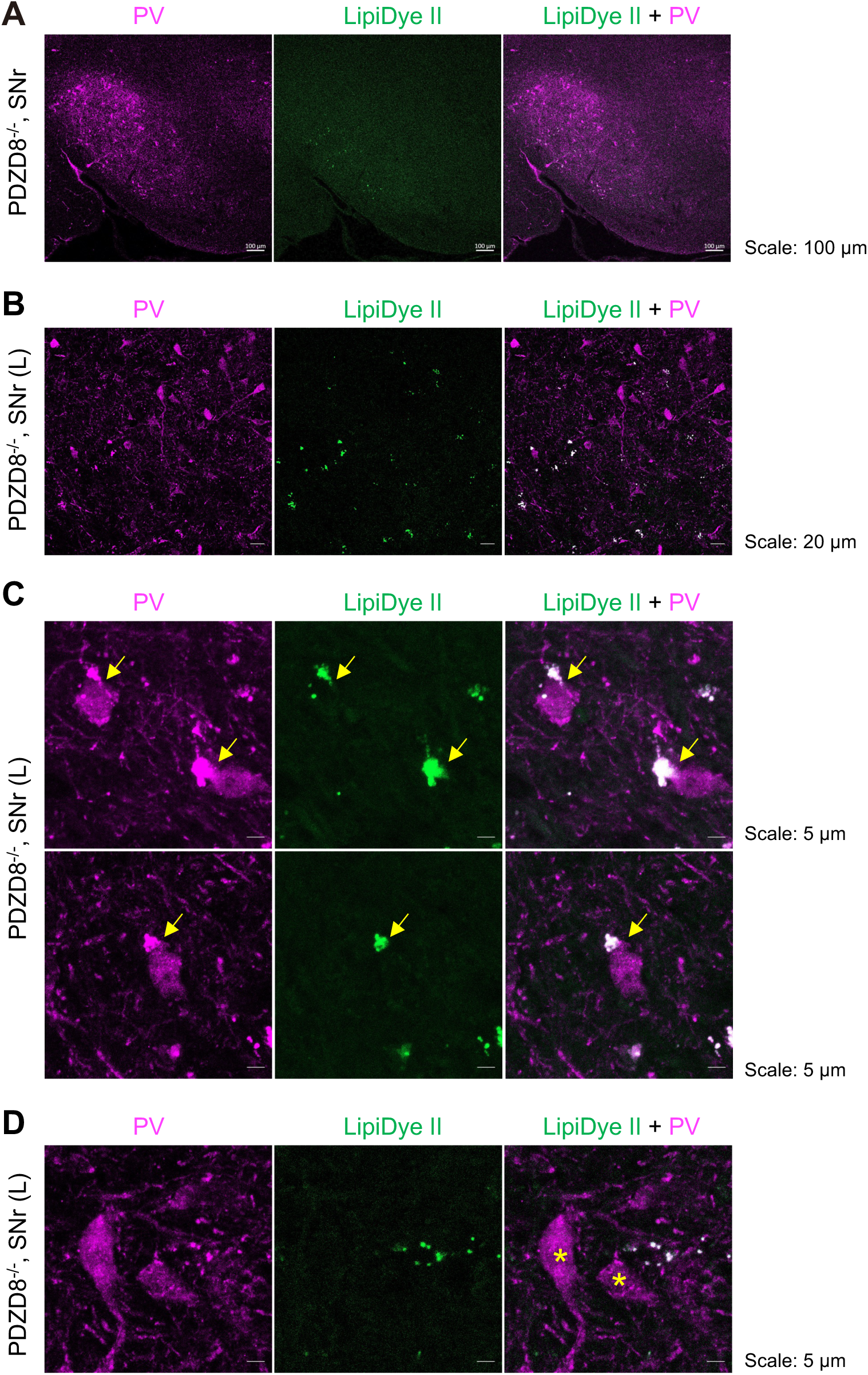
Lipofuscin in SNr in PDZD8^−/−^ mice. The same experiments as in Figure 4F with other mice. Confocal images of LipiDye II (green) and PV (magenta) in PV neurons containing lipofuscin deposits in lateral regions of caudal SNr in PDZD8^−/−^ mice. (A) Caudal SNr corrresponding to atlas #86. Scale bars, 100 µm. (B) Lateral region in (A). Scale bars, 20 µm. (C) PV neurons containing lipofuscin deposits in (B). PV neurons with LipiDye II-positive structure (yellow arrows) are indicated. Scale bars, 5 µm. (D) PV neurons without lipofuscin deposits in (B). PV neurons without LipiDye II-positive structure (asterisks) are indicated. Scale bars, 5 µm.

**Supplementary figure 5.**
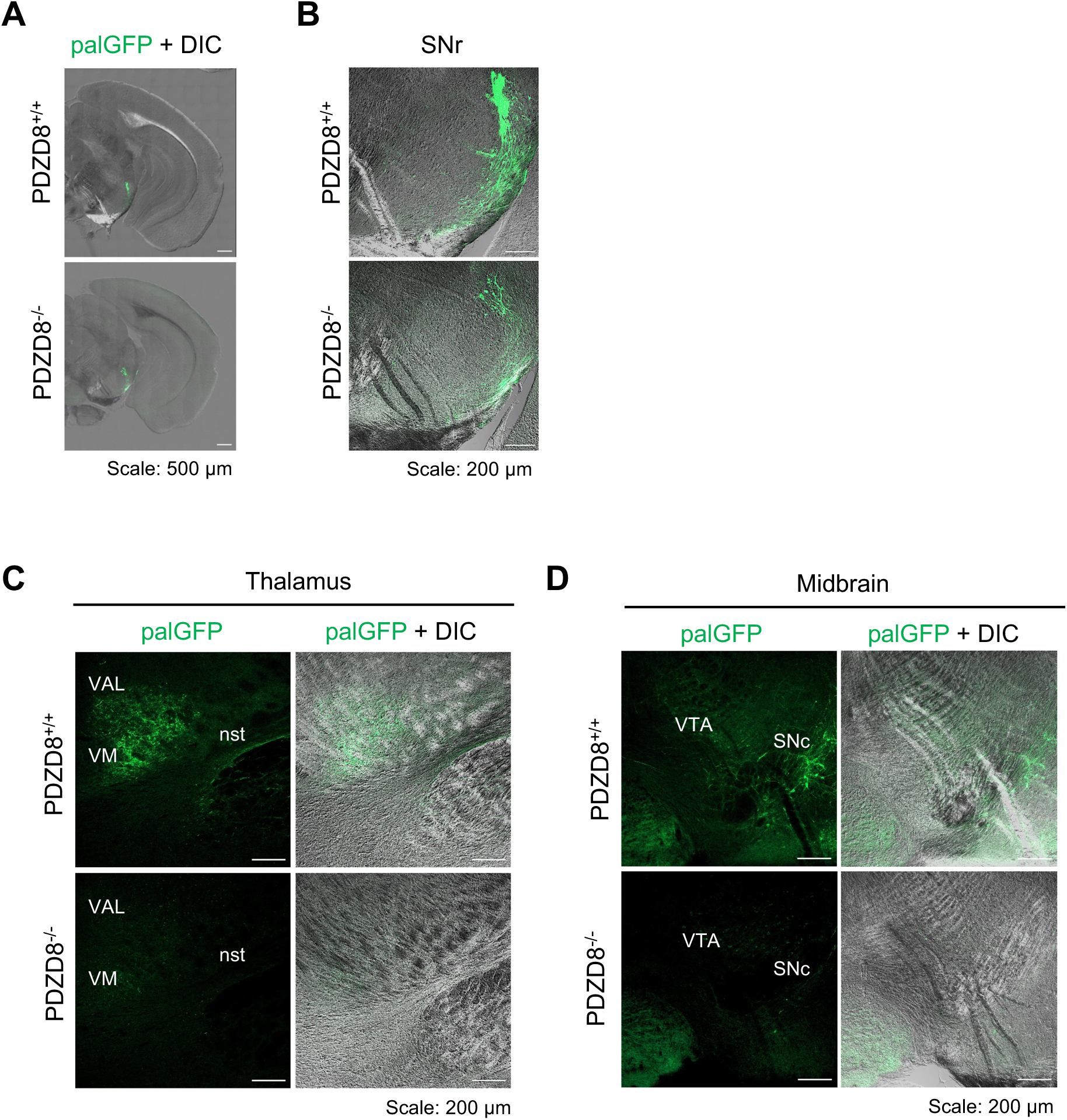
Decreased projection from SNr in PDZD8^−/−^ mice. The same experiments as in Figure 5 with another mouse pair. (A, B) Confocal images showing palmitoylated GFP (palGFP) injection sites in SNr merged with DIC images. Tiling images of cerebral hemisphere (A) and sections focusing on SNr (B). Scale bars, 500 µm (A); 200 µm (B). (C) Confocal palGFP images merged with DIC of thalamic regions showing SNr projection targets. PalGFP signals indicate SNr projections to the ventral anterior-lateral complex of the thalamus (VAL), ventromedial thalamus (VM), and the nigrostriatal tract (nst). Scale bars, 200 µm. (D) Confocal and merged DIC images of midbrain regions showing SNr projections to the ventral tegmental area (VTA) and substantia nigra pars compacta (SNc). Scale bars, 200 µm.

**Supplementary figure 6.**
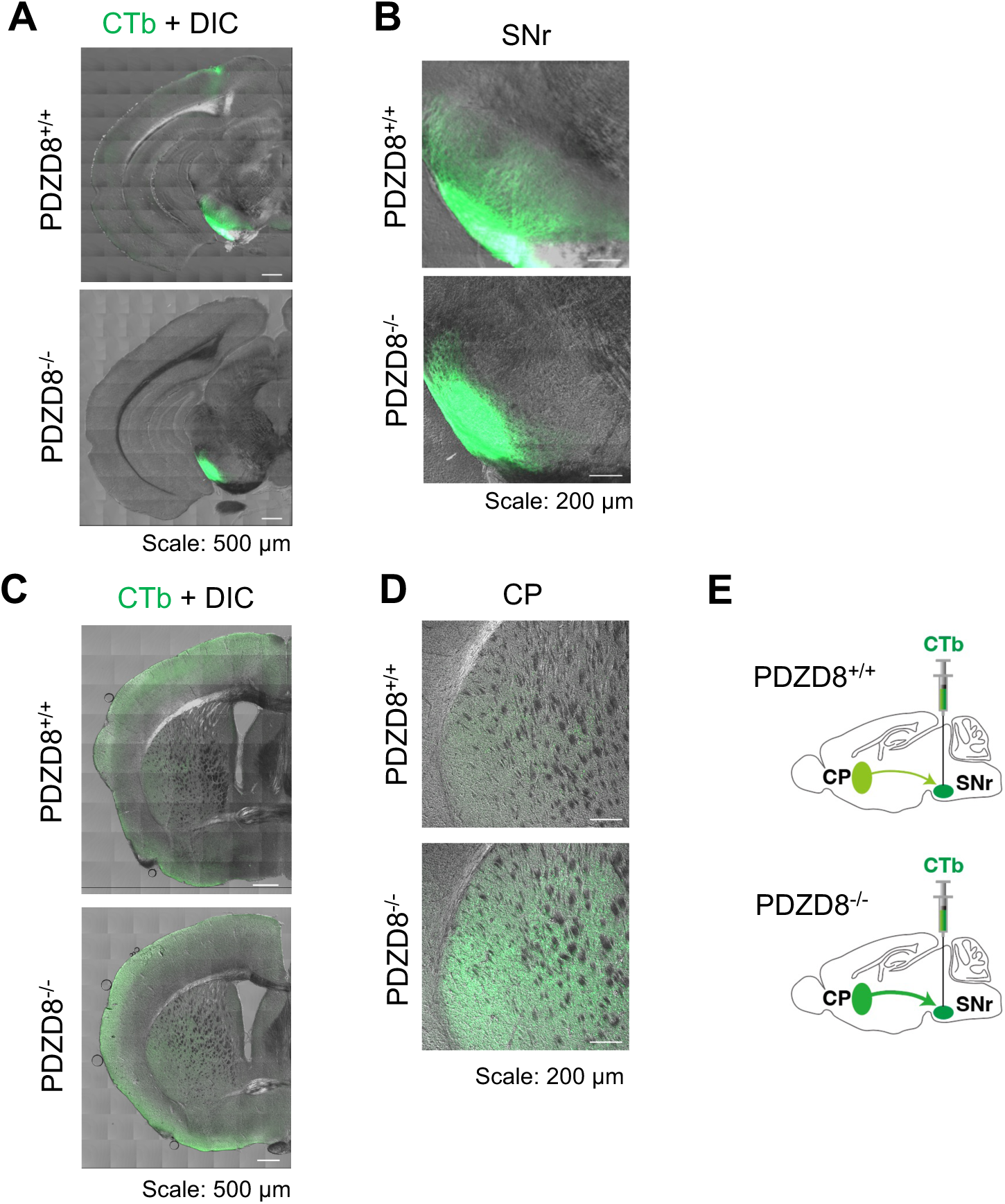
Increased striatal projections to the SNr in PDZD8^−/−^ mice. (A, B) Confocal images of Alexa488-CTb injected to SNr merged with DIC images. Tiling images of cerebral hemisphere (A) and images focused on SNr (B). Scale bars, 500 µm (A) and 200 µm (B). (C, D) Confocal images of Alexa488-CTb merged with DIC images of brain region containing caudate-putamen (CP) in striatum, indicating projection sources. Tiling images of cerebral hemisphere (C) and images focused on CP (D). Scale bars, 500 µm (C) and 200 µm (D).

**Supplementary table 1.** Information of mouse age and sex. The age and sex of mice used in this study are summarized.

**Information of mouse age and sex**
| **Main figure** | **Age (month)** | **Sex (M/F)** |
| --- | --- | --- |
| 1A | 7 | F |
| 1B | 7 | F |
| 1C | 7 | F |
| 1D | 9 | F |
| 2A | 4 | M |
| 2B | 4 | M |
| 2C | 4 | M |
| 2D | 4 | M |
| 3A-D | 9 | M |
| 3E, F | 6 | M |
| 4B, C | 4 | M |
| 4D, E | 9 | M |
| 4F | 4 | M |
| 4F | 7 | F |
| 5A-E | 4 | M |
| 6A-F | 4 | F |
| 6H | 4 | M |
| 7A, B | 8 | F |
| 7C | 4 | M |
| **Supplement** | **Age (month)** | **M/F** |
| S1A,B | 6 | F |
| S1C | 4 | M |
| S2A | 6 | F |
| S2B | 5 | F |
| S3A, B | 6 | M |
| S3C | 9 | M |
| S4A-D | 4 | M |
| S5A-D | 4 | M |
| S6A-D, 6F | 4 | M |
| SmovA, B | 9 | M |

**Supplementary table 2.**
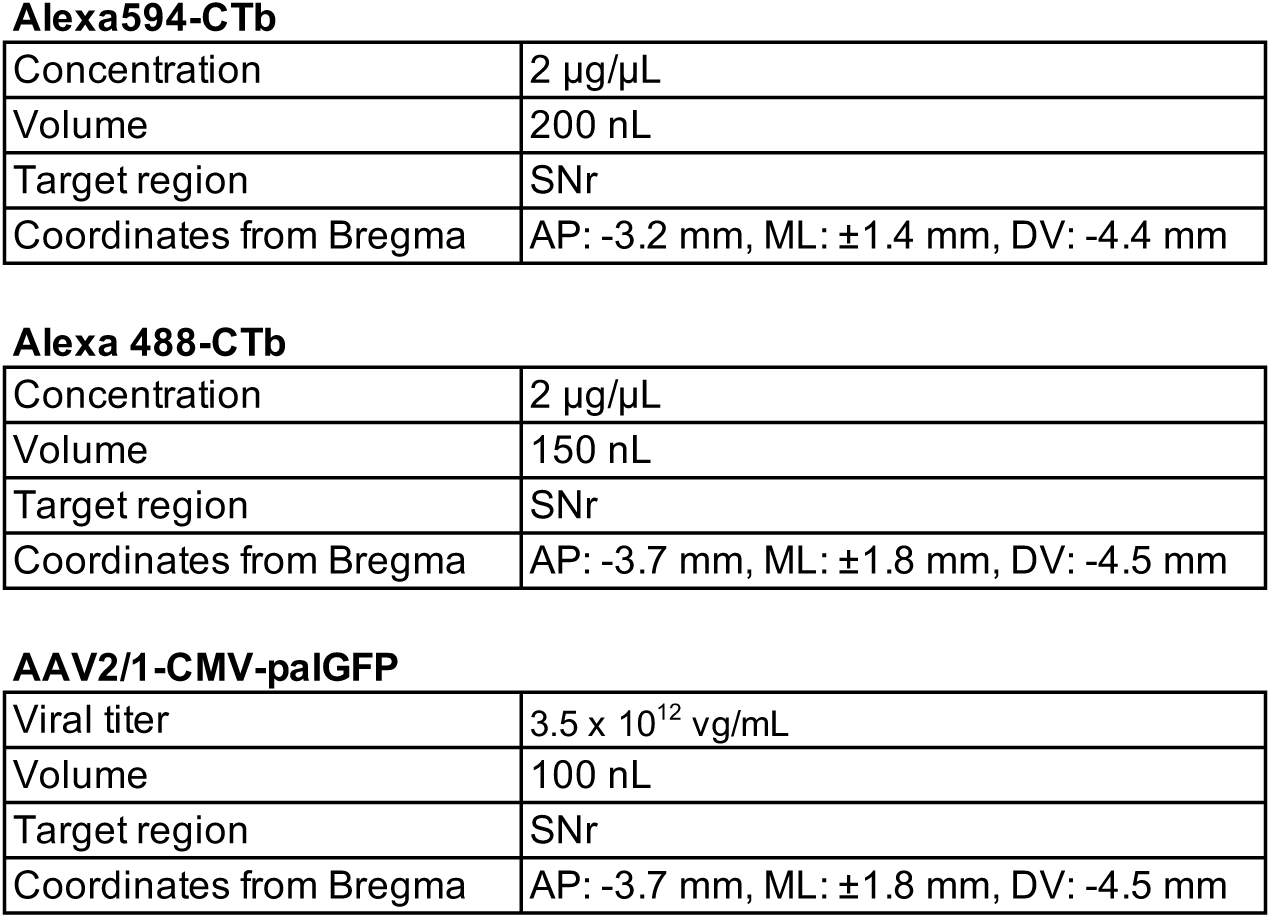
Information of neuronal tracer injection. Concentration, volume, target region, and coordinates from Bregma for Alexa594-CTb, Alexa488-CTb, and AAV2/1-CMV-palGFP were summarized.

**Alexa594-CTb**
|  |  |
| --- | --- |
| Concentration | 2 µg/µL |
| Volume | 200 nL |
| Target region | SNr |
| Coordinates from Bregma | AP: -3.2 mm, ML: ±1.4 mm, DV: -4.4 mm |
| **Alexa 488-CTb** |  |
| Concentration | 2 µg/µL |
| Volume | 150 nL |
| Target region | SNr |
| Coordinates from Bregma | AP: -3.7 mm, ML: ±1.8 mm, DV: -4.5 mm |
| **AAV2/1-CMV-paIGFP** |  |
| Viral titer | 3.5 x 10<sup>12</sup> vg/mL |
| Volume | 100 nL |
| Target region | SNr |
| Coordinates from Bregma | AP: -3.7 mm, ML: ±1.8 mm, DV: -4.5 mm |

**Supplementary movies. 3D images of lipid deposits in SNr in PDZD8^−/−^ mice.** (A, B) 3D movies of LipiDye II (green) and GFAP (magenta) in medial (A) and lateral (B) regions of SNr in PDZD8^−/−^ mice, which are corresponding to Figure 4B and C, respectively. Scale bars, 10 µm.

## Notes

### Competing Interest Statement

The authors have declared no competing interest.

### Summary of Updates

In the revised version, the causative mechanism was added, that is, Lipofuscin deposits in PV neurons in SNr of PDZD8 KO mice leads to dysfunction in basal ganglia circuitry.

